# Spermine inhibits pathogen-associated molecular pattern (PAMP) ROS and Ca^2+^ burst and reshapes the transcriptional landscape of PAMP-triggered immunity in *Arabidopsis thaliana*

**DOI:** 10.1101/2022.05.05.490826

**Authors:** Chi Zhang, Kostadin E. Atanasov, Rubén Alcázar

## Abstract

Polyamines are small polycationic amines which levels increase during defense. Previous studies support the contribution of the polyamine spermine (Spm) to the establishment of the hypersensitive response (HR) during incompatible plant-pathogen interactions. However, the potential contribution of Spm to other layers of defense, and pathogen-associated molecular pattern (PAMP)-triggered immunity (PTI) in particular, was not completely established. Here we compared the contribution of Spm and putrescine (Put) to early and late PTI responses. We find that Put and Spm show opposite effects on PAMP-elicited reactive oxygen species (ROS) production, with Put increasing whereas Spm lowering flg22-stimulated ROS burst. Through genetic and pharmacological approaches, we find that the inhibitory effect of Spm on flg22-elicited ROS is independent of polyamine oxidation and EDS1 (ENHANCED DISEASE SUSCEPTIBILITY 1), PAD4 (PHYTOALEXIN DEFICIENT 4), salicylic acid and NPR1 (NONEXPRESSER OF PR GENES 1) defense components but resembles chemical inhibition of RBOHD (RESPIRATORY BURST OXIDASE HOMOLOG D) function. Remarkably, Spm can also suppress ROS elicited by FLS2-independent but RBOHD-dependent pathways, thus pointing to compromised RBOHD function. Consistent with this, we find that Spm dampens flg22-stimulated cytosolic Ca^2+^ influx necessary for RBOHD function and reshapes the transcriptional landscape of PTI and defense responses against *Pseudomonas syringae* pv. *tomato* DC3000. Overall, we provide molecular evidence for the differential contribution of Put and Spm to PTI with an impact on plant defense.

## INTRODUCTION

The most abundant polyamines in plants are the diamine putrescine (Put), triamine spermidine (Spd) and tetramine spermine (Spm). The contents of polyamines are increased in response to abiotic and biotic stresses. Polyamine levels are regulated through tight control of their biosynthesis, oxidation by polyamine oxidases (PAO) and copper-containing amine oxidases (CuAO), conjugation to hydroxycinnamic acids, acylation and transport (Cona et al., 2006; Alcázar et al., 2010; Tiburcio et al., 2014; Zeiss et al., 2021). Increasing evidence support the contribution of polyamines to biotic stress resistance, although their effects on defense signaling are not completely established (Walters, 2003; Tiburcio et al., 2014; Seifi and Shelp, 2019). We recently reported that Put is synthesized in response to systemic acquired resistance (SAR)-inducing bacteria and this polyamine triggers local salicylic acid (SA) accumulation and systemic responses contributing to SAR establishment and defense against *P. syringae* (Liu et al., 2020). Studies in Arabidopsis and tobacco also indicate that Spm enhances resistance against Cauliflower mosaic virus, *P. viridiflaba, P. syringae, Hyaloperonospora arabidopsidis, Verticillium dahliae* and *Botrytis cinerea*, among other pathogenic microorganisms. In most cases, Spm responses were found to be dependent on polyamine oxidation (Marina et al., 2008; Mitsuya et al., 2009; Moschou et al., 2009; Sagor et al., 2012; Marco et al., 2014; Mo et al., 2015). Spm also activates the protein kinases SIPK (salicylic acid (SA)-induced protein kinase) and WIPK (wound-induced protein kinase) in tobacco (Takahashi et al., 2003), as well as mitogen-activated protein kinases (Zhang and Klessig, 1997; Seo et al., 2007) leading to the expression of a number of hypersensitive response (HR) marker genes in a ROS (reactive oxygen species) and Ca^2+^-dependent but SA-independent manner (Takahashi et al., 2004). Overall, the data suggest the contribution of Spm to defense through potentiation of HR. However, the potential contribution of Spm to other layers of defense, and pathogen-associated molecular pattern (PAMP)-triggered immunity (PTI) in particular, has not been established. *Pseudomonas syringae* produces the small molecule phevamine A, a modified form of Spd that suppresses the potentiation effect of this polyamine on flagellin-stimulated ROS burst (O’Neill et al., 2018). Therefore, polyamine analogs can be used by pathogens to subvert PTI responses.

Plants have two layers of pathogen recognition (Dodds and Rathjen, 2010). The first layer is initiated upon perception of PAMPs by pattern recognition receptors (PRRs), which leads to PTI. A second intracellular layer relies on nucleotide-binding domain and leucine-rich repeat-containing receptor (NLR) proteins, which directly or indirectly recognize virulence effectors and induce effector-triggered immunity (ETI). The Arabidopsis leucine-rich repeat receptor kinase FLS2 (FLAGELLIN SENSITIVE 2) recognizes the bacterial flagellin (Gómez-Gómez et al., 2000; Zipfel, 2014). Binding of the immunogenic flagellin peptide (flg22) initiates a number of downstream responses. One of the earliest signaling events after PAMP recognition is a rapid increase in cytosolic Ca^2+^ levels ([Ca^2+^]_cyt_), ROS generation, and activation of MAPKs and Ca^2+^-dependent protein kinases (CPKs) ultimately leading to transcriptional and metabolic reprogramming (Boller and Felix, 2009; Segonzac and Zipfel, 2011). Ca^2+^ is a ubiquitous second messenger which signal specificity is explained by the duration, amplitude, frequency and spatial distribution of the Ca^2+^ burst. Specific Ca^2+^ signatures are decoded by Ca^2+^-binding proteins that translate this information into changes in the phosphorylation status of proteins and transcriptional responses (Dodd et al., 2010). ROS has been proposed to act as antimicrobial, facilitate cell wall modifications and act in local and systemic defense signaling (Lamb and Dixon, 1997; Suzuki et al., 2011; Nathan and Cunningham-Bussel, 2013). ROS are generated by different enzymatic complexes including Class III peroxidases, oxalate oxidases, lipoxygenases, quinone reductases, amine oxidases including CuAO and PAO, and NADPH oxidases (Cona et al., 2006; Miller et al., 2010). ROS production during PTI is predominantly dependent on the NADPH oxidase RBOHD (RESPIRATORY BURST OXIDASE HOMOLOG D) (Nühse et al., 2007; Zhang et al., 2007). In general, RBOH proteins transfer electrons from cytosolic NADPH or NADH to apoplastic oxygen, producing superoxide (O^2-^) which can be converted to hydrogen peroxide (H_2_O_2_) by superoxide dismutases (Marino et al., 2012; Suzuki et al., 2012). RBOHs have Ca^2+^-binding EF-hand motifs in their N-terminal region that bind Ca^2+^. Indeed, Ca^2+^-binding is important for RBOHD function since treatment with Ca^2+^ chelators and point mutations in EF-hand motifs compromise PAMP-triggered ROS production (Kadota et al., 2004; Ogasawara et al., 2008; Ranf et al., 2011; Segonzac and Zipfel, 2011; Kimura et al., 2012; Kadota et al., 2014). ROS produced by RBOHD activity also induces Ca^2+^ influx, thus suggesting a positive feedback regulation that boosts ROS production (Ranf et al., 2011). Other mechanisms of RBOHD regulation involve phosphorylation at different sites by the protein kinase BIK1 (BOTRYTIS-INDUCED KINASE1) and CPKs upon PAMP perception (Boudsocq et al., 2010; Suzuki et al., 2011; Marino et al., 2012; Dubiella et al., 2013; Kadota et al., 2014; Li et al., 2014). RBOHD phoshorylation by BIK1 is independent of calcium-based regulatory mechanisms, but Ca^2+^ is required for ultimate PAMP-triggered RBOHD activation (Kadota et al., 2014). In addition, RBOHs are also regulated by binding of small GTPases, 14-3-3 proteins and phosphatidic acid (Morel et al., 2004; Elmayan et al., 2007; Wong et al., 2007; Zhang et al., 2009). The many regulatory mechanisms, as well as the broad range of functions on stress and development of RBOH family members, suggest that they act as molecular hubs mediating ROS-signaling.

In this work, we investigated the effect of polyamines on PAMP-elicited ROS burst, which is one of the earliest PTI responses. By focusing on flg22-elicitation of PTI, we find that Spm strongly inhibits flg22-mediated ROS production. In contrast, pretreatment with Put enhanced flg22-triggered ROS. Through genetic and pharmacological approaches, we provide evidence that the inhibitory effect of Spm on flg22-ROS burst does not require polyamine oxidation, is independent of EDS1 (ENHANCED DISEASE SUSCEPTIBILITY 1), PAD4 (PHYTOALEXIN DEFICIENT 4), SA and NPR1 (NONEXPRESSER OF PR GENES 1) and cannot be ameliorated by Put treatment. Inhibition ROS production by Spm is also observed in response to other FLS2-independent but RBOHD-dependent ROS-inducing agents. Spm mimics the effect of Ca^2+^ chelators and Ca^2+^ channel blockers that compromise RBOHD function. In agreement, we find that Spm dampens flg22-triggered Ca^2+^ influx required for RBOHD function and reshapes the transcriptional landscape of PTI with an impact on disease resistance against *P. syringae*.

## RESULTS

### Effect of Put and Spm on flg22-triggered ROS burst

To study the effect of different polyamines on PTI, we first analyzed the contribution of Put and Spm to flg22-triggered ROS burst in Arabidopsis. ROS production was measured in wild-type plants treated with flg22 (1 μM) supplemented with different concentrations of Put and Spm (50, 100, 200 and 400 μM) or mock **(Figure 1)**. Cotreatments of flg22 with Put produced no evident changes on flg22-triggered ROS burst (**Figure 1A**). In contrast, preincubation with Put (100 μM) 24 h before flg22 elicitation triggered higher ROS production compared to mock pretreatment (**Figure 1B**). This effect was absent at lower or higher Put concentrations. In contrast, 100 μM and higher concentrations of Spm strongly inhibited flg22-triggered ROS production (**Figure 1A and 1B**). The cotreatment of flg22 with Put and Spm also led to ROS burst inhibition (**Figure S1**). The inhibitory effect of Spm on flg22-triggered ROS was also evidenced in defense mutants *eds1-2, pad4-1, sid2-1* and *npr1-1* **(Figure S2)**, thus pointing to an EDS1/PAD4, SA and NPR1 independent response.

**Figure 1.**
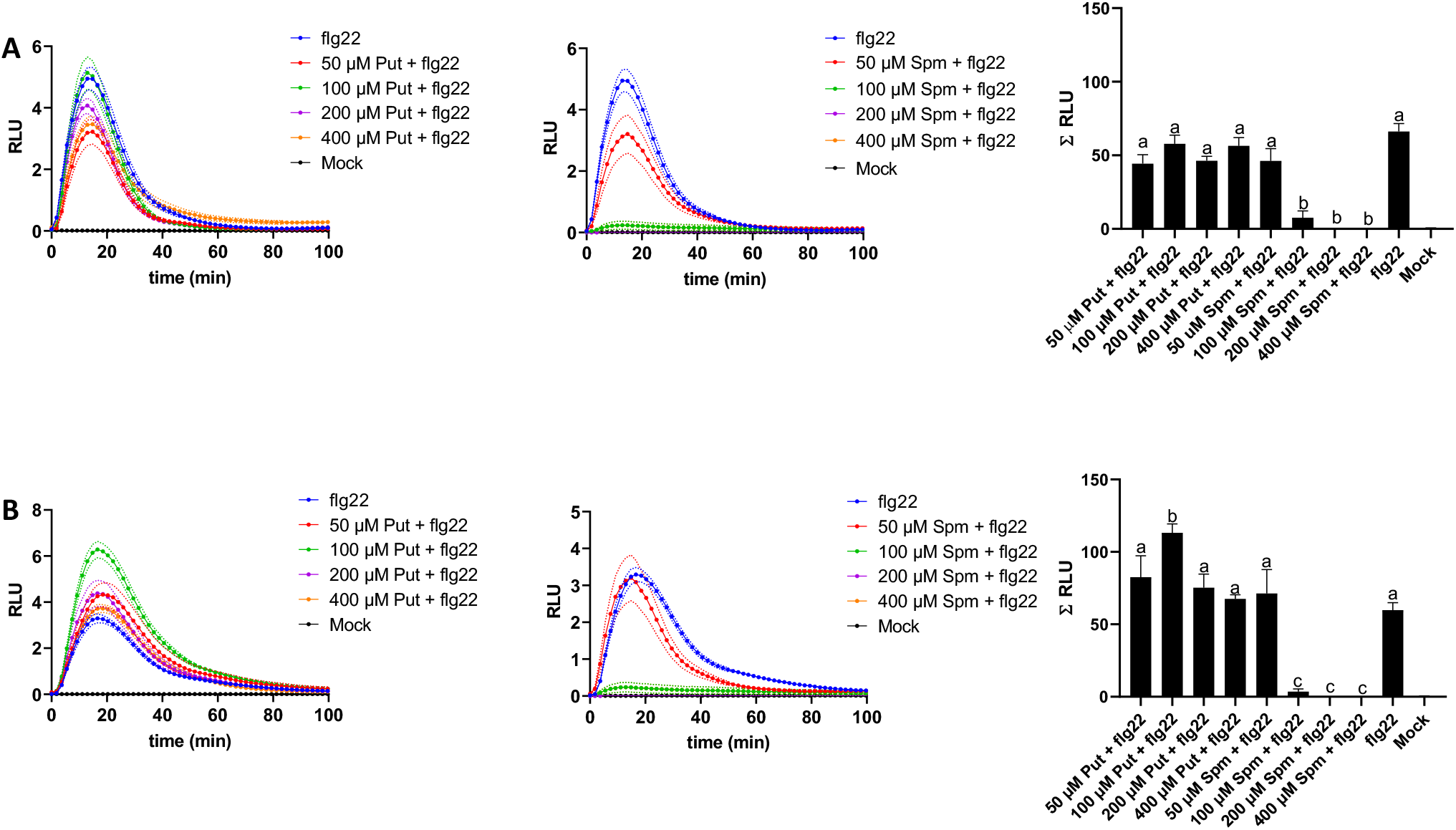
Effect of Put and Spm on flg22-elicited ROS burst. **(A)** Leaf discs from fully expanded leaves of 5-week-old wild-type (Col-0) plants were treated with flg22 (1 μM) and Put or Spm at indicated concentrations (50 μM to 400 μM). **(B)** Leaf discs were preincubated with Put or Spm at indicated concentrations 24 h before flg22 (1 μM) elicitation. Values represent the mean ± S.E. from at least twelve replicates per treatment and are expressed in relative light units (RLU). Letters indicate values that are significantly different according to Tukey’s HSD test at *P* < 0.05.

Incubation with the individual polyamines (Put or Spm) at different concentrations produced much lower but sustained ROS, which showed a plateau between four to eight hours of 100 μM Put or 100 μM Spm treatments (**Figure S3**). The polyamine-triggered ROS production kinetics did not overlap with the flg22-elicited ROS production response. Interestingly, Spm-triggered ROS production was abrogated in *rbohd* but not *rbohf* mutants (**Figure S4**). The results indicated that apoplastic ROS stimulated by Spm is RBOHD-dependent.

### Flg22 triggered ROS burst in *adc1, adc2* and *spms* mutants

To further study the effect of Put and Spm on flg22-triggered ROS burst, we used *arginine decarboxylase 1 (adc1-3)* and *arginine decarboxylase 2 (adc2-4)* mutants compromised in Put biosynthesis, and *spermine synthase (spms)* mutant deficient in Spm biosynthesis (**Figure 2**). The *adc1-3* and *adc2-4* mutants showed similar flg22-ROS production compared to the wild-type (**Figure 2A**). In contrast, flg22-triggered ROS levels were significantly higher in *spms* than in wild-type plants (**Figure 2B**). The data was consistent with an inhibitory effect of Spm on flg22-ROS burst (**Figure 1**). We concluded that Put and Spm have opposite effects on flg22-triggered ROS production, with Put increasing whereas Spm lowering the amplitude of the ROS response.

**Figure 2.**
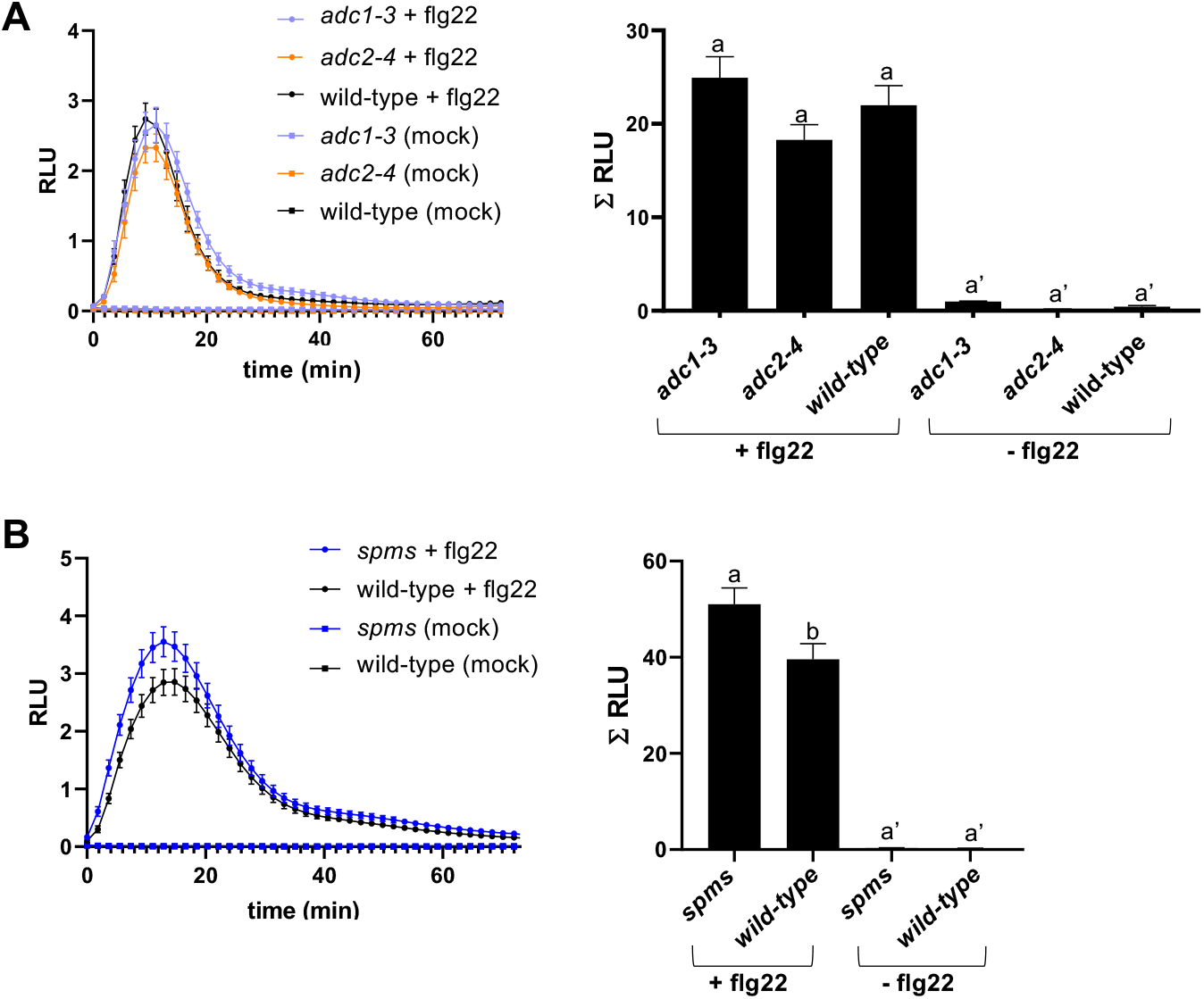
Flg22-stimulated ROS burst in **(A)** *adc1-3, adc2-4* and **(B)** *spms* mutants. Leaf discs from fully expanded leaves of 5-week-old wild-type (Col-0) and mutants were treated with flg22 (1 μM) or mock (water). Values represent the mean ± S.E. from at least twelve replicates per treatment and are expressed in relative light units (RLU). Letters indicate values that are significantly different according to Tukey’s HSD test at *P* < 0.05.

### Effect of Put and Spm on flg22-triggered *Pst* DC3000 disease resistance

Pre-treatment of wild-type plants with flg22 induces resistance to *Pseudomonas syringae* pv. *tomato* DC3000 (*Pst* DC3000) (Zipfel et al., 2004). Given the opposite effect of Put and Spm on flg22-triggered ROS burst, we determined *Pst* DC3000 disease resistance phenotypes in wild-type plants and *eds1-2* mutant treated with flg22 (1 μM), Put (100 μM), Spm (100 μM), combinations (100 μM Put + 1 μM flg22) and (100 μM Spm + 1 μM flg22), and mock (**Figure 3**). Wild-type plants preinfiltrated with (Spm + flg22) supported higher bacteria growth than plants pretreated with flg22 or (Put + flg22) (**Figure 3A**). These results were consistent with the observed inhibition of flg22 ROS burst caused by Spm, which could partly compromise flg22-elicited defenses (**Figures 1 and 2**). In contrast, pretreatment of wild-type plants with the individual polyamines (100 μM) did not lead to significant changes on *Pst* DC3000 disease resistance compared to mock (**Figure 3A**). The data indicated that Spm is not an unspecific suppressor of defense responses. No differences were observed in bacteria growth between the different treatments in the *eds1-2* mutant (**Figure 3B**).

**Figure 3.**
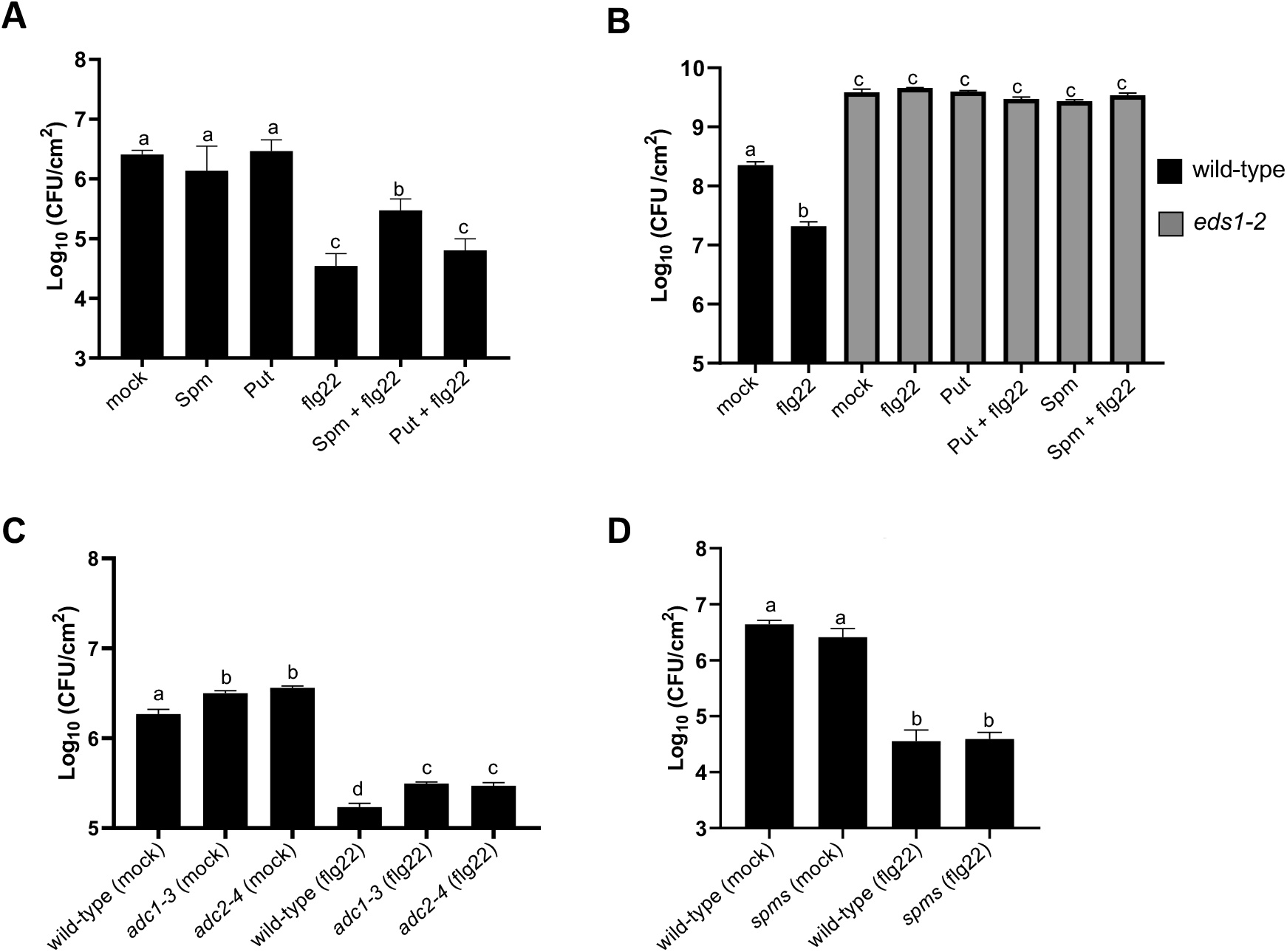
Analysis of *Pst* DC3000 disease resistance in **(A)** wild-type plants and **(B)** *eds1-2* mutant preinfiltrated with mock (10 mM MgCl_2_), Put (100 μM), Spm (100 μM), flg22 (1 μM) and combinations (Spm + flg22, Put + flg22). **(C, D)** Disease resistance phenotypes to *Pst* DC3000 in **(C)** *adc1-3, adc2-4* and **(D)** *spms* mutants preinfiltrated with flg22 (1 μM) or mock (10 mM MgCl_2_). Preinfiltrations were performed 24 h before *Pst* DC3000 inoculation. In all treatments, fully expanded leaves of five-week-old *Arabidopsis* plants were infiltrated with *Pst* DC3000 (OD_600_ nm= 0.005). Bacterial numbers were assessed at 72 h post-inoculation and expressed as colony forming units (CFU) per cm^2^ leaf area. Values are the mean from at least eight biological replicates ±SD. Letters indicate values that are significantly different according to Tukey’s HSD test at *P* < 0.05.

Despite we did not detect increased resistance to *Pst* DC3000 at 100 μM Put, higher concentrations of Put (200 μM to 500 μM) caused lower bacteria growth in the wild-type (**Figure S5**), which otherwise did not correlate with the amplitude of the flg22-ROS burst (**Figure 1B**). The *adc1-3* and *adc2-4* mutants were more susceptible to infection by *Pst* DC3000 than wild-type plants in both flg22 and mock pre-inoculated plants (**Figure 3C**). The data supported a positive contribution of Put to defense independently of FLS2 signaling. Infiltration of the *spms* mutant with *Pst* DC3000 led to similar bacteria growth to the wild-type. In addition, no significant differences in *Pst* DC3000 growth were observed between wild-type and *spms* plants pretreated with flg22 (**Figure 3D**). This indicated that the enhanced flg22-ROS burst in *spms* (**Figure 2B**) was not associated with higher disease resistance. Due to the striking effect of Spm on flg22-triggered ROS inhibition, we focused on this polyamine for further analyses.

### Spm oxidation is not required for flg22-triggered ROS burst inhibition

To determine whether polyamine oxidation by polyamine oxidases (PAO) or copper-containing amine oxidases (CuAO) was necessary for the inhibitory effect of Spm on flg22-ROS burst, we determined ROS production in response to flg22 (1 μM) and (100 μM Spm + 1 μM flg22) in *CuAO* mutants (*atao1, cuao1, cuao2, cuaoα1, cuaoα*2, *cuaoδ, cuaoε*1, *cuaoε*2, *cuaoγ*2) and *PAO* mutants (*pao1, pao2, pao3, pao4* and *pao5*). No significant differences were detected in flg22-elicited ROS production between *cuao, pao* and wild-type plants (**Figure S6A**). In addition, amine oxidase mutations or treatment with the amine oxidase inhibitor 2-bromoethylamine (BEA) did not rescue the effect of Spm on flg22-ROS burst inhibition (**Figures S6B and S7**). We concluded that polyamine oxidation was dispensable for the inhibitory effect of Spm on flg22-elicited ROS production.

### Spm inhibits RBOHD-dependent ROS production

To further study the inhibitory effect of Spm on flg22-triggered ROS burst, we performed a pharmacological approach using wild-type plants treated with the NADPH oxidase inhibitor diphenyleneiodonium (DPI), ROS scavengers dimethylthiourea (DMTU) and reduced L-glutathione (GSH), Ca^2+^ chelator EGTA, Ca^2+^ channel blocker lanthanum chloride (LaCl_3_) and the protein synthesis inhibitor cycloheximide (CHX). We also included the actin depolymerization inhibitor latrunculin B (Lat B) that strongly reduces flg22-induced FLS2 internalization and the brassinosteroid (BR) biosynthesis inhibitor brassinazole (BRZ) in the analyses (Albrecht et al., 2012; Belkhadir et al., 2012) (**Figures S7 and S8**).

As expected, flg22-triggered ROS production was absent in the treatments that compromise RBOHD function (DPI, EGTA and LaCl_3_) and ROS production (DMTU and GSH). The Flg22-triggered ROS inhibition by these chemicals resembled Spm treatment (**Figure S7**). In contrast, treatments with Lat B or BRZ did not rescue flg22-triggered ROS production in the presence of Spm (**Figures S7 and S8A**). We concluded that the inhibitory effect of Spm on flg22-ROS burst was not due to FLS2 internalization or *de novo* BR biosynthesis (Albrecht et al., 2012; Belkhadir et al., 2012). Interestingly, treatment with CHX produced high ROS levels that were absent in (CHX + Spm) treatments (**Figures S7 and S8B**). The data indicated that the inhibitory effect of Spm on flg22-ROS burst does not require *de novo* protein biosynthesis. Furthermore, CHX-triggered ROS production was compromised in *rbohd* but not *fls2*, which supported its dependence on RBOHD independently of FLS2 (**Figures S8B and S8C**). We hypothesized that Spm inhibits flg22-triggered ROS production through its ROS scavenging capacity and/or inhibition of RBOHD function.

### Analysis of the ROS scavenging and cell death trigger capacity of Spm

To investigate the potential ROS-scavenging capacity of Spm in plants, we performed 3,3’ diaminobenzidine (DAB) staining in wild-type and *rbohd* leaves infiltrated with Spm (100 μM), methyl viologen (MV) (100 μM), Spm (100 μM) + MV (100 μM) and mock (**Figure S9A**). Infiltration with MV led to uniform DAB precipitates in wild-type and *rbohd* leaves, indicative of high H_2_O_2_ production independently of RBOHD. In contrast, infiltration with Spm (100 μM) did not lead to evident DAB staining and resembled mock treatment. Co-infiltration of MV + Spm only partly alleviated the presence of DAB precipitates, which otherwise were still evident in wild-type and *rbohd* leaves (**Figure S9A**). MV, Spm or MV+Spm did not induce cell death in any of the genotypes tested, as revealed by trypan blue staining (**Figure S9B**). The data indicated that Spm (100 μM) only partly reduces ROS levels in Arabidopsis leaves and, at this concentration, is not a cell death trigger.

### Effect of Spm on flg22-induced calcium influx

PAMPs induce a rapid transient increase in cytosolic Ca^2+^ ([Ca^2+^]_cyt_) by influx of Ca^2+^ from the extracellular environment or internal stores (McAinsh and Pittman 2009). This Ca^2+^ burst operates upstream of many PAMP-elicited responses and is necessary for RBOHD function (Blume et al., 2000; Boller and Felix, 2009; Ranf et al., 2011; Segonzac et al., 2011). We used the apoaquorein bioluminescent Ca^2+^ sensor to measure steady-state levels and dynamics of [Ca^2+^]_cyt_ in response to flg22, Spm and combinations in the wild-type background (**Figure 4**). Flg22 treatment triggered a rapid Ca^2+^ influx which was partly inhibited by Spm. Spm treatment also triggered increases in [Ca^2+^]_cyt_, although the amplitude of the Ca^2+^ response was broader than flg22 or (Spm + flg22) treatments (**Figure 4A**). We then analyzed the effect of flg22 or Spm pre-treatment on successive flg22/Spm-triggered Ca^2+^ burst. Pretreatment with Spm 30 min before flg22 elicitation led to much lower [Ca^2+^]_cyt_ increases (**Figure 4B**). On the other hand, flg22 pretreatment also compromised Spm-triggered Ca^2+^ burst (**Figure 4B**). The data indicated that Spm triggers cytosolic Ca^2+^ influx and dampens the Ca^2+^ burst mediated by flg22. We reason that the inhibitory effect of Spm on flg22-stimulated Ca^2+^ burst might compromise RBOHD activation and thus, flg22-triggered ROS production.

**Figure 4.**
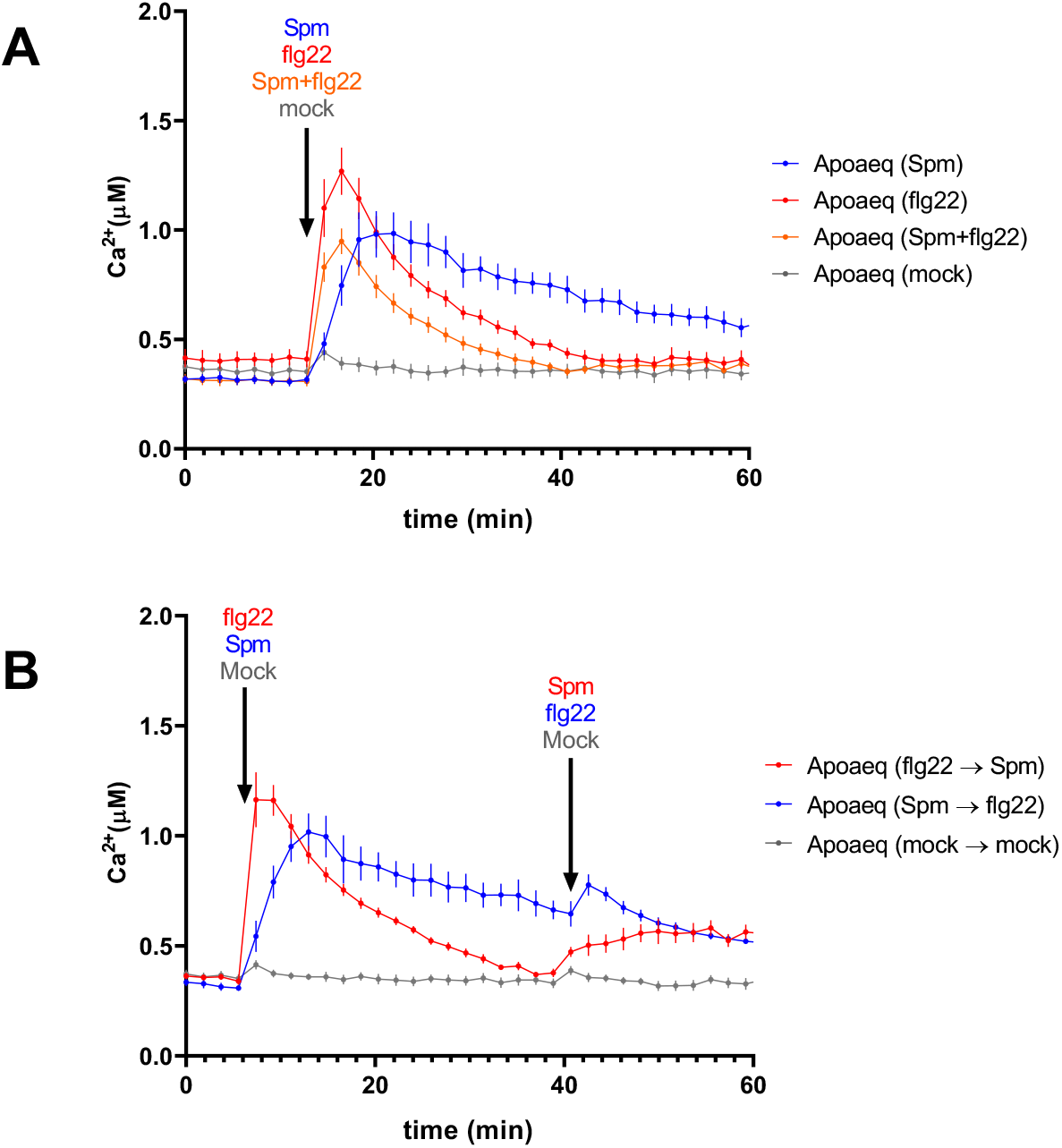
Flg22- and Spm-induced Ca^2+^ signatures.**(A)** Leaf discs of apoaequorin-expressing line (Knight et al., 1991) were treated with 1 μM flg22, 100 μM Spm or mock (water) and luminescence monitored over time. **(B)** Effect of flg22 or Spm pre-treatments on subsequent Ca^2+^ burst elicited by Spm or flg22, respectively. Values represent the mean ± S.E. from at least twelve replicates per treatment and are expressed in relative light units (RLU).

### Comparative RNA-seq gene expression analyses of wild-type plants treated with Put and Spm

Given the contrasting effects of Put and Spm on flg22-triggered ROS burst, we performed gene expression analyses (RNA-seq) to identify differences in the transcriptional responses triggered by these two polyamines at 24 h of treatment (**Figure 5; Tables S1.1 to S1.8**). A total of 554 and 368 genes were significantly deregulated (fold-change ≥ 2.0 and Bonferroni corrected p-value ≤ 0.05) after 100 μM Put or 100 μM Spm treatments, respectively (**Figure 5A; Tables S1.1 and S1.2**). Closer inspection of Put responsive genes identified an overrepresentation of enzymes (84 genes) mostly downregulated and involved in the metabolism of sugars, lipids, amino acids and nitrogenous bases, among others (**Figure 5B; Table S1.1**). Put-deregulated genes also included 81 downregulated transcription factors (TFs) enriched in members of the APETALA 2/ETHYLENE RESPONSE ELEMENT BINDING PROTEINS (AP2/EREBP) family, 35 upregulated ribosomal or rRNA processing proteins, 20 genes encoding cell wall biogenesis or modification enzymes (most upregulated) and other 20 RING/U-box family proteins (most downregulated) (**Figure 5; Table S1.1**). Among the genes only upregulated by Put, we found a significant enrichment in gene ontology (GO) terms related to translation, whereas Put-only downregulated genes were related to stress (**Figure 5; Table S1.6**). Low correlation was found (r^2^ =0.4149) between Put and Spm treatments in the set of Put-only deregulated genes (**Figure 5**).

**Figure 5.**
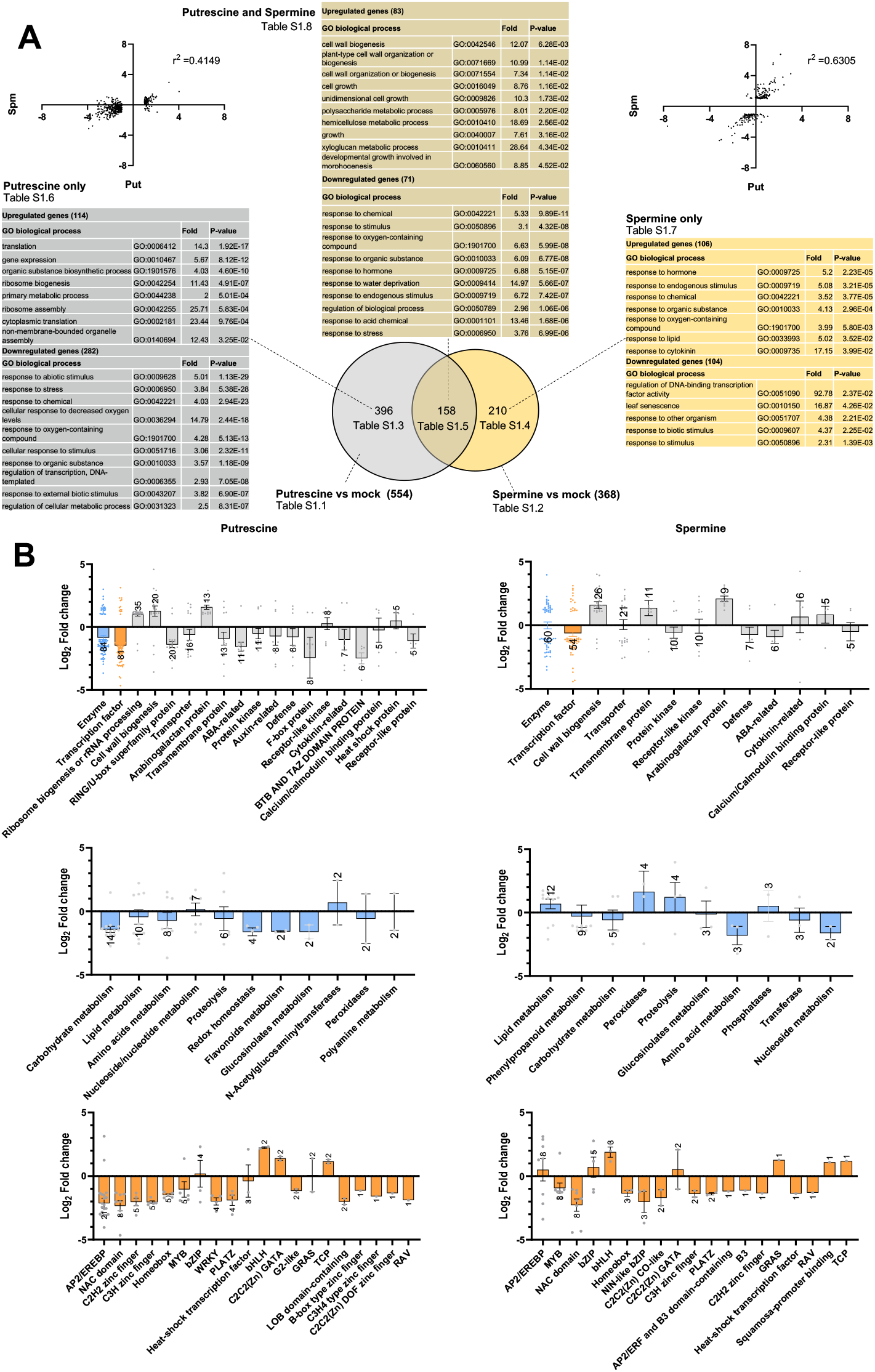
**(A)** Venn diagram, gene ontology (GO) and expression correlation analyses of genes significantly deregulated (fold-change ≥2; Bonferroni corrected p-value ≤0.05) in response to Put (100 μM) and Spm (100 μM) at 24 h of treatment in the wild-type. **(B)** Molecular functions, main enzymatic activities and TF families of genes differentially expressed in Put and Spm treatments. Bars indicate the mean expression ± S.E. Number of genes within each category are indicated on top of the bars. A complete list of genes is found in Tables S1.1 to S1.8.

Spm-responsive genes were also enriched in enzymes (60 genes) involved in lipid metabolism (upregulated), phenylpropanoid and sugar metabolism (downregulated) (**Figure 5; Table S1.2**). Spm-deregulated genes included 54 TFs with an equal representation of AP2/EREBP (most upregulated), MYB and NAC domain containing families (downregulated) (**Figure 5; Table S1.2**). Other 26 Spm-upregulated genes were related to cell wall biogenesis, of which half of them (13 genes) were common with Put (**Figure 5; Tables S1.1 and S1.2**). The set of Spm-only upregulated genes was enriched in hormone, lipid and cytokinin responses. Spm-only downregulated genes included GO terms related to the regulation of TF activity, senescence and biotic responses (**Figure 5 and Table S1.7**). Spm-only deregulated genes also showed low correlation (r^2^ =0.6305) between Spm and Put treatments (**Figure 5**). The data was consistent with a differential regulation of gene expression by these two polyamines. Indeed, only 158 genes were commonly deregulated by Put and Spm (**Table S1.5**). Upregulated genes within this expression sector were mainly related to cell wall biogenesis and organization (**Figure 5 and Table S1.8**). Overall, Put showed a transcriptional effect on ribosome biogenesis which was not evidenced in the Spm treatment. The polyamines also showed different effects on the expression of primary metabolism genes and TF gene families. However, both polyamines enhanced the expression of genes encoding enzymes that modify the composition and assembly of the cell wall carbohydrate fraction.

### Effect of Spm and Put on flg22-elicited transcriptional responses

To further investigate the effect of Put and Spm on flg22-triggered responses, we determined the changes in expression at 24 h of flg22 (1 μM), Spm (100 μM) + flg22 (1 μM), Put (100 μM) + flg22 (1 μM) and mock treatments in wild-type plants. The RNA-seq data was used for principal component analysis (PCA) and hierarchical clustering analysis (HCA) (**Figure 6; Tables S2.1 to S2.19; Tables S3.1 to S3.19**). The PC1 of the PCA explained 32 % of total variance and mainly differentiated between flg22-treated and flg22-untreated samples (**Figure 6A**). Flg22 treatment also discriminated between the two major clades of the HCA analysis (**Figure 6B**). The PC2, (16.1 % of total variance), revealed the differences in expression due to the different polyamines (**Figure 6A**), which also grouped into separate subclades of the HCA analysis (**Figure 6B**).

**Figure 6.**
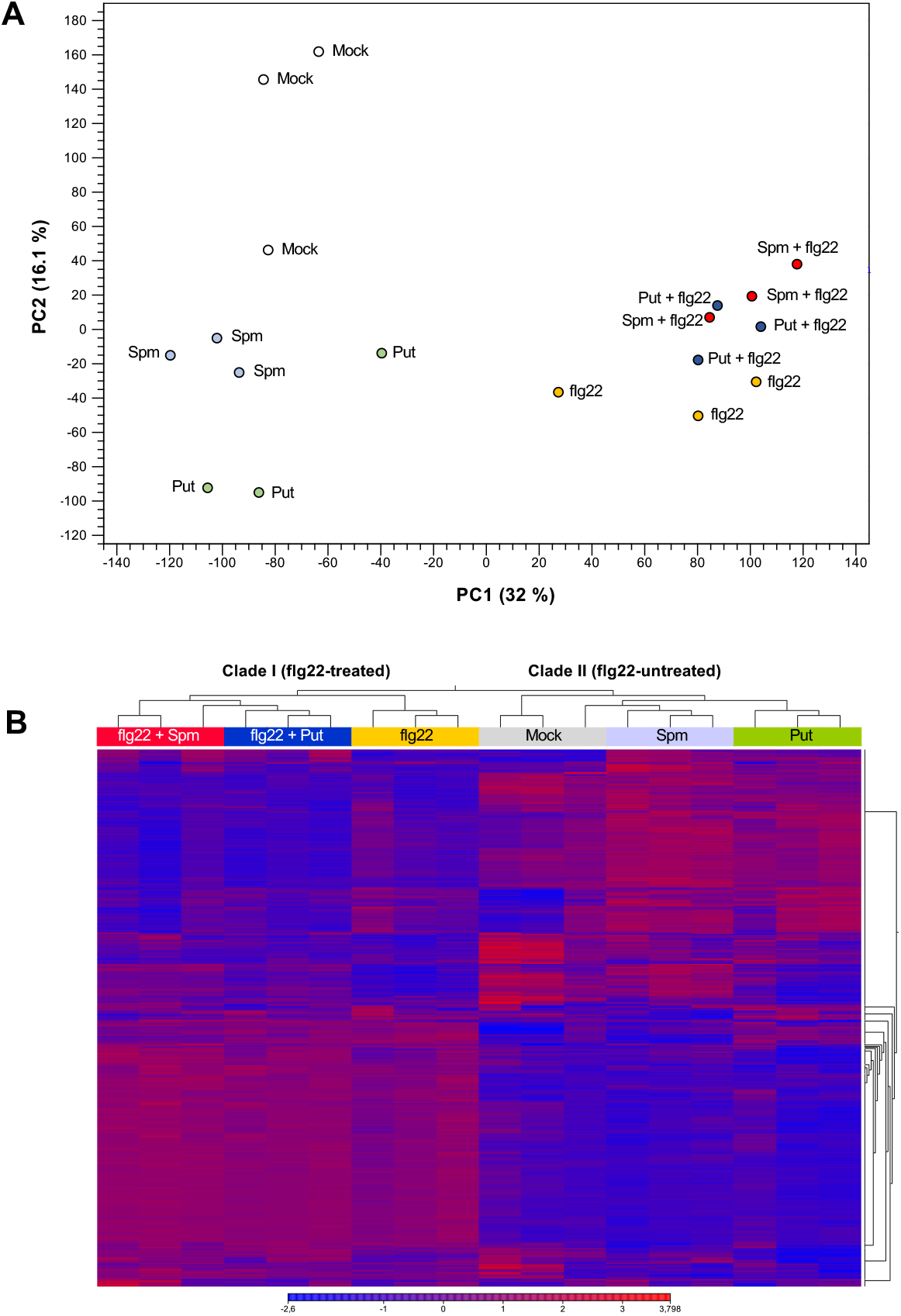
**(A)** Principal component analysis (PCA) and **(B)** Hierarchical clustering analysis (HCA) of RNA-seq gene expression data obtained from 5-week-old wild-type plants treated with Put (100 μM), Spm (100 μM), flg22 (1 μM), Put (100 μM) + flg22 (1 μM), Spm (100 μM) + flg22 (1 μM) and mock (water) for 24 h. Each treatment was performed in three biological replicates.

### Comparative gene expression analysis of wild-type plants treated with (Spm + flg22) and flg22

Compared to mock, flg22 triggered the deregulation of 1415 genes (**Figure 7; Table S2.1**). Most flg22-responsive genes were related to defense but also included genes involved in ribosome biogenesis (**Tables S2.1 and S2.2**). Out of the 1415 flg22-responsive genes, 462 (33 %) were not significantly deregulated by flg22 in the presence of Spm (**Figures 7 and S10A; Table S2.3**). This set of genes was enriched in ribosomal proteins (55 genes) and GO terms related to translation (**Figure S11; Tables S2.3 and S2.4**). The flg22-only sector also included genes involved in phenylpropanoid biosynthesis (upregulated), and downregulated genes related to carbon starvation, biosynthesis of amino acids, glucosinolates, trehalose and fatty acid metabolism, among others (**Figure S11; Table S2.5**). The data suggested that Spm dampens the expression of a subsector of flg22-responsive genes involved in ribosomal biogenesis and stress metabolism adaptation.

**Figure 7.**
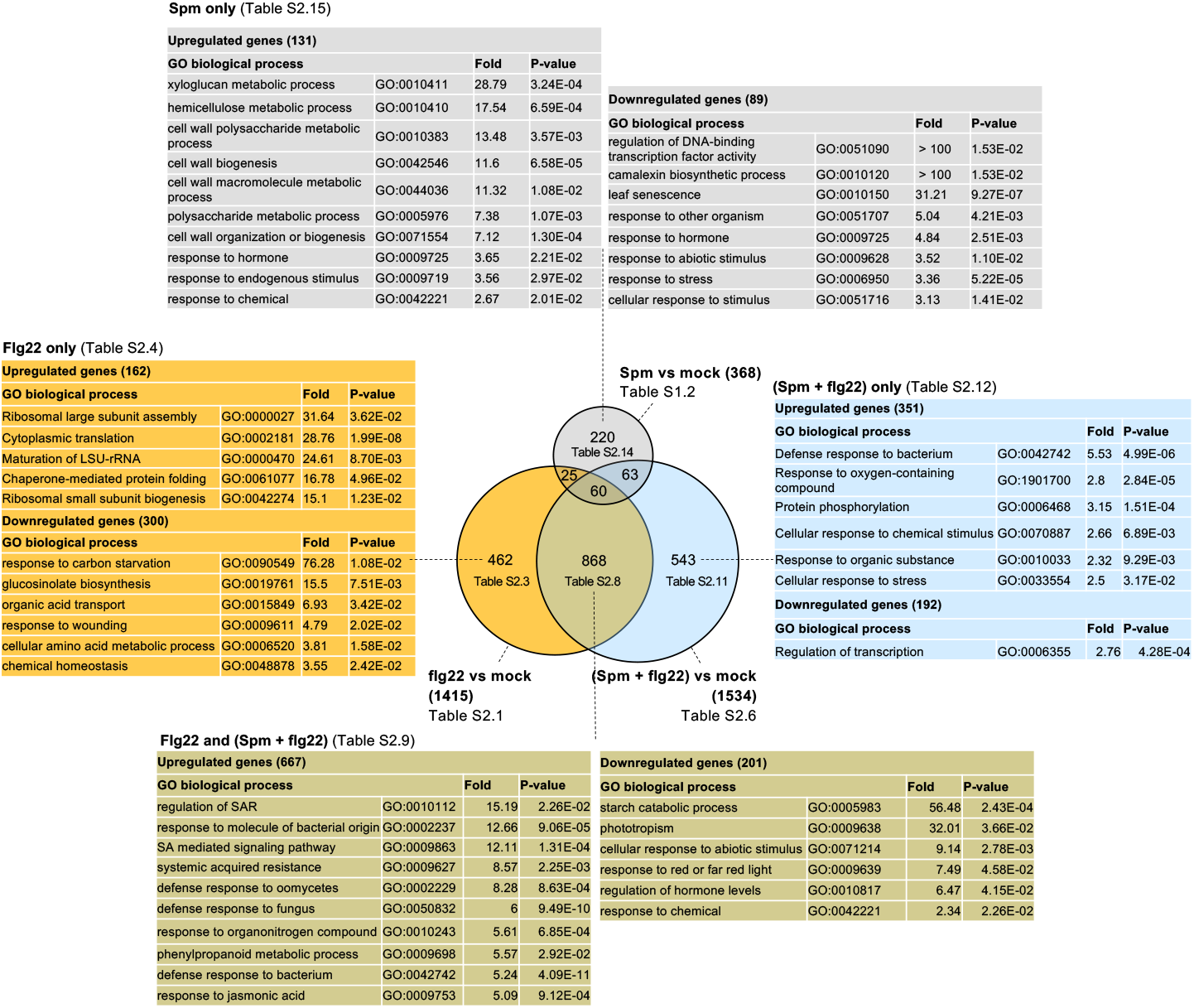
Venn diagram and gene ontology (GO) analyses of genes differentially expressed (fold change ≥2; Bonferroni corrected p-value ≤0.05) in response to flg22 (1 μM), Spm (100 μM) and Spm (100 μM) + flg22 (1 μM). The fold-enrichment (Fold) and Bonferroni-corrected p-values for each GO term are indicated. Genes within each category are listed in Tables S2.1 to S2.19.

Treatment of wild-type plants with (Spm + flg22) triggered the deregulation of 1534 genes (**Figure 7; Table S2.6**), of which only 868 (**Figure S10B**; **Table S2.8**) and 63 (**Table S2.17**) were common with flg22 and Spm treatments, respectively. The flg22 and (Spm + flg22) common genes were enriched in GO categories related to SAR, defense, SA and JA responses, phenylpropanoid metabolism (all upregulated), plant hormone biosynthesis and starch degradation (downregulated) among other biological functions (**Figures 7 and S12, Tables S2.9 and S2.10**). Most abundant molecular functions included enzymes (246 genes), transporters (60 genes), transcription factors (55 genes), ribosomal proteins and rRNA processing enzymes (47 genes) (**Figure S12 and Table S2.8**). The mean expression of upregulated genes within this sector was slightly but significantly higher in (Spm + flg22) than flg22 treatment (**Figure S10B**). The (Spm + flg22)-only sector included 543 genes which biological functions were related to defense responses to bacterium, ROS responses, protein phosphorylation, flavonoid biosynthesis (all upregulated); and transcription regulation (downregulated), among others (**Figures 7, S10C and S13; Tables S2.11 to S2.13**). This set of (Spm + flg22)-only genes was used to determine correlation coefficients between flg22 and (Spm + flg22) treatments. We found that both treatments were correlated (r^2^ = 0.9036), despite the difference in the number of genes assigned to each sector (**Figure S10E**). This is probably due to the threshold criteria used to identify differentially expressed genes (fold change ≥ 2 and Bonferroni corrected p-value ≤ 0.05) that overestimates the differences between treatments and the observation that Spm increases the expression of many flg22-responsive genes (**Figure S10B; Table S2.8**). The correlation between flg22 and (Spm+flg22) treatments in genes within the flg22-only sector was lower (r^2^ = 0.8056) (**Figure S10E**). Genes exclusively deregulated by Spm were significantly enriched in GO terms related to cell wall biogenesis (upregulated) (**Figures 7, S10D, and S14; Tables S2.14 and S2.15**), which suggested that polyamines contribute to cell wall reinforcement and modifications during defense. In this case, no strong correlations were detected between Spm and (Spm + flg22) treatments (r^2^ =0.5353) (**Figure S10E**). We concluded that Spm boosts the expression of a significant number of flg22-responsive genes related to defense and cell wall modifications, while it decreases the expression of ribosomal proteins and genes related to metabolism adaptation during defense activation (**Figure 7**).

### Comparative gene expression analysis of wild-type plants treated with (Put + flg22) and flg22

In the Put treatment, out of the 1415 flg22-responsive genes, 259 (18.3 %) were not significantly deregulated by flg22 in the presence of Put (flg22-only in **Figure 8 and S15A; Table S3.1**). However, flg22 and (Put + flg22) treatments were highly correlated (r^2^ = 0.9439) (**Figure S15E**), which suggested that the significance threshold overestimated the qualitative differences between treatments. Indeed, differences were more quantitative than qualitative given that Put dampened the expression of flg22-responsive genes in this sector (**Figure S15A; Table S3.1**). Flg22-only genes were also enriched in ribosomal proteins (30 genes) (**Figures 8 and S16; Tables S3.1 and S3.2**). However, these accounted for almost half of the ribosomal genes upregulated by Spm treatment (55 genes) (**Table S2.3**). The “Put and flg22” (91 genes) and “Put, flg22 and (Put + flg22)” (163 genes) expression sectors were also enriched in GO terms related to translation (**Figure 8 and Tables S3.15 to S3.18**), consistent with the effect of Put on upregulation of genes encoding ribosomal proteins and rRNA processing enzymes **(Figure 5; Table S1.1)**. Flg22-only upregulated genes also included enzymes (59) enriched in the phenylpropanoid pathway that are also upregulated in the common flg22 and (Put+flg22) gene expression sector (**Figures S16 and S17; Tables S3.1 to S3.3 and S3.6 to S3.8**).

**Figure 8.**
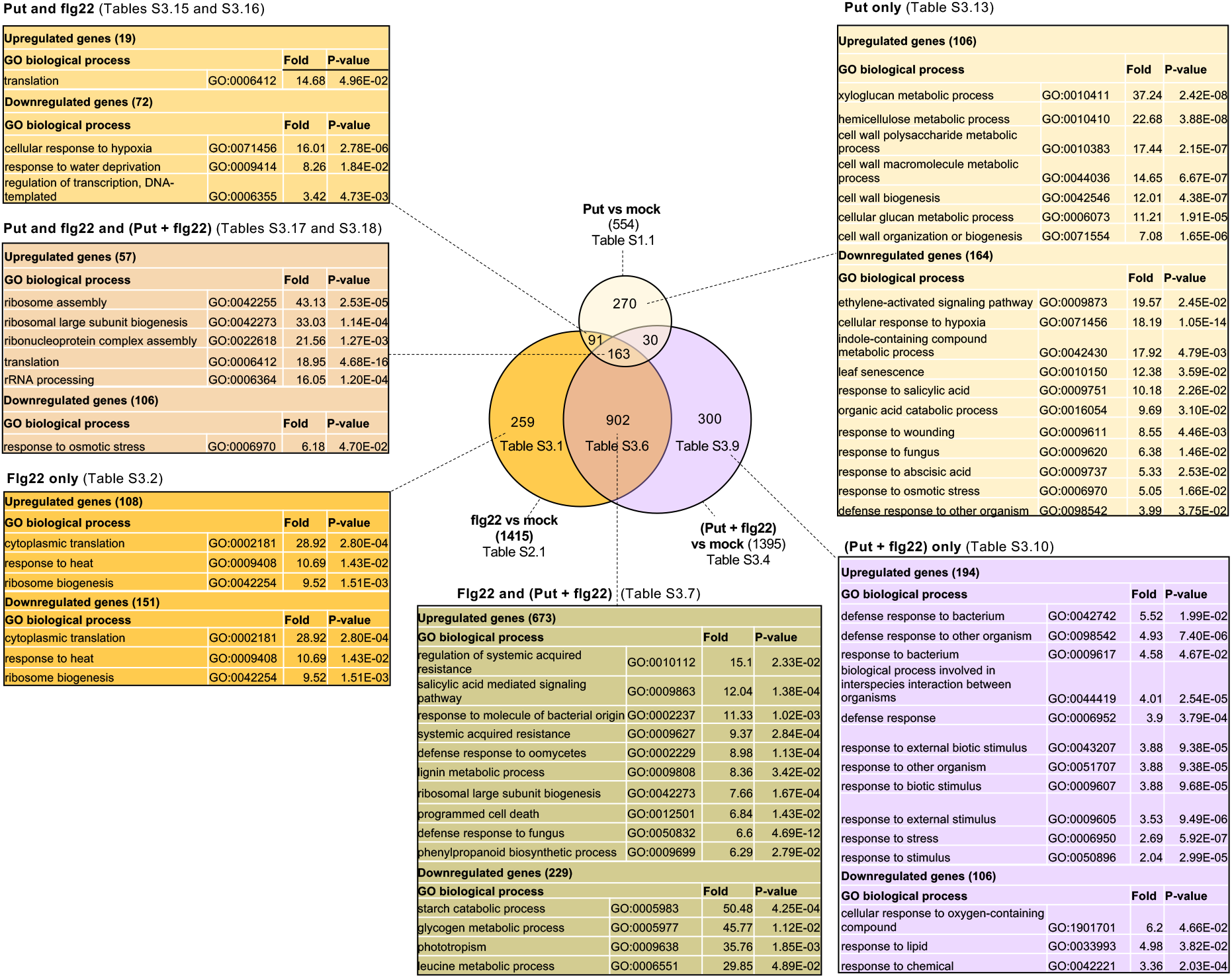
Venn diagram and gene ontology (GO) analyses of genes differentially expressed (fold change ≥2; Bonferroni corrected p-value ≤0.05) in response to flg22 (1 μM), Put (100 μM) and Put (100 μM) + flg22 (1 μM). The fold-enrichment (Fold) and Bonferroni-corrected p-values for each GO term are indicated. Genes within each category are listed in Tables S3.1 to S3.19.

Treatment of wild-type plants with (Put + flg22) triggered the deregulation of 1395 genes (**Figure 8; Table S3.4**), of which 902 (**Figure S15B**; **Table S3.6**) and 30 (**Table S3.19**) were common with flg22 and Put treatments, respectively. The flg22 and (Put + flg22) common expression sector was enriched in GO categories related to SAR, SA signaling, defense, lignin metabolism, ribosome biogenesis, phenylpropanoid metabolism (all upregulated) and starch catabolism (downregulated) among other biological functions (**Figures 8 and S17, Tables S3.6 to S3.8**). Most abundant molecular functions included enzymes (252 genes), transporters (67 genes), transcription factors (53 genes) and others including 26 ribosomal proteins (**Figure S17 and Table S3.6**). Similarly to (Spm + flg22) treatment, (Put + flg22) also led to a slighter but still significant increase in the mean expression of flg22 responsive genes (**Figure S15B**). The (Put + flg22)-only sector included 300 genes mainly enriched in defense responses to bacterium (upregulated), response to oxygen-containing compound and response to lipids (downregulated) (**Figures 8, S15C and S18; Tables S3.9 to S3.11**). The expression of genes within this sector was also correlated with flg22 treatment (r^2^ = 0.8757), which accounted for quantitative rather than qualitative differences between the sectors (**Figure S15E**). Like in the case of Spm, genes only deregulated by Put were mostly enriched in cell wall biogenesis functions (upregulated) and showed low correlation with other treatments (**Figures 8, S15D, S15E and S20; Tables S3.12 to S3.14**).

Collectively, the data indicated that Spm, and less markedly Put, compromise flg22-induced transcriptional upregulation of genes encoding ribosomal proteins, enhance the expression of a subsector of flg22-responsive genes related to defense and trigger the upregulation of cell wall biogenesis and modification enzymes which are not deregulated by flg22, thus reshaping the PAMP-induced transcriptional responses.

## DISCUSSION

Spm and its precursors Put and Spd show opposite effects on the stimulation of flg22-elicited ROS production and defense elicitation. While Spm inhibits flg22-triggered ROS burst (**Figures 1A and 1B**), Put (**Figure 1B**) and Spd (**Figure S20**) (O’Neill et al., 2018) potentiate ROS production. Importantly, the inhibitory effect of Spm does not require its oxidation (**Figures S6B and S7**) and cannot be compensated by Put (**Figure S1**), which apoplastic levels are increased in response to *Pst* DC3000 inoculation (Liu et al., 2019). Upon flg22 binding, FLS2 forms a complex with BAK1 (BRI1-ASSOCIATED KINASE1), an LRR-receptor kinase that also serves as coreceptor of BRI1 (BRASSINOSTEROID INSENSITIVE1) involved in BR signaling (Chinchilla et al., 2009; Postel et al., 2010; Schulze et al., 2010; Fradin et al., 2011; Schwessinger et al., 2011). Brassinolides (BL) have also been shown to inhibit FLS2 signaling, including flg22-elicited ROS burst in Arabidopsis, downstream or independently of the FLS2-BAK1 complex or through competition of BAK1 recruitment by FLS2 and the BR receptor BRI1 (Albrecht et al., 2012; Belkhadir et al., 2012). However, treatment with the BL biosynthesis inhibitor brassinazole (BRZ) did not rescue the inhibitory effect of Spm on flg22-triggered ROS production (**Figure S8A**). Also, compromised FLS2 internalization by treatment with Lat B did not rescue Spm-triggered ROS inhibition (**Figure S7**). In addition to flg22-triggered ROS burst, Spm also inhibited CHX-induced ROS production, which was RBOHD-dependent but FLS2-independent (**Figures S8B and S8C**). The data suggested that Spm inhibition of flg22-elicited ROS was not due to impaired FLS2 signaling. Rather, Spm mimicked the inhibitory effect of the NADPH oxidase inhibitor DPI, Ca^2+^ chelator EGTA and Ca^2+^ channel blocker lanthanum chloride (LaCl_3_) on flg22-triggered ROS production (**Figure S7**). Flg22-elicited ROS is mainly contributed by RBOHD, which activation requires Ca^2+^ binding to N-terminal EF-hand motifs (Kadota et al., 2004; Ogasawara et al., 2008; Ranf et al., 2011; Segonzac and Zipfel, 2011; Kimura et al., 2012; Kadota et al., 2014). Flg22 treatment triggered a rapid [Ca^2+^]_cyt_ increase that was significantly dampened by Spm (**Figure 4**). On the other hand, Spm also triggered Ca^2+^ influx but the Ca^2+^ signature lasted longer than with flg22 treatment. Because Ca^2+^ burst stimulated by Spm was also inhibited by flg22 pretreatment, we concluded that flg22 and Spm produce different Ca^2+^ signatures that are reciprocally inhibitory (**Figure 4**). The suppressive effect of Spm on flg22-triggered Ca^2+^ influx might explain the inhibition of flg22-elicited ROS production by this polyamine.

(Rocha et al., 2020) reported that Spm was necessary for mucilage production during appressoria morphogenesis in the blast fungus *Magnaporthe oryzae*, by buffering oxidative stress in the ER lumen and preventing the unfolded protein response. The antioxidant properties of Spm have been documented and involve the conversion of the polyamine into different adducts, including Spm dialdehyde, in the presence of hydroxyl radicals (Ha et al., 1998). In this case, the potential ROS scavenging capacity of Spm is unlikely to explain its inhibitory effect on flg22-elicited ROS production. We found that Spm (100 μM) only shows partial antioxidant capacity in Arabidopsis leaves infiltrated with MV, a ROS inducing agent that transfers electrons from photosystem I to molecular oxygen (**Figure S9**). In addition, Spm but also its precursor Spd act as ROS scavengers through seemingly similar mechanisms (Khan et al., 1992). However, these two polyamines show opposite effects on flg22-triggered ROS stimulation (**Figures 1 and S20**) (O’Neill et al., 2018). Furthermore, we found that thermospermine (tSpm, 100 μM), an isomer of Spm, does not inhibit flg22-triggered ROS production (**Figure S20**).

Polyamines are known to affect ion transport across membranes through intricate mechanisms which are also dependent on polyamine charge (Pottosin and Shabala, 2014). Spm can induce membrane depolarization or hyperpolarization, depending on the supplied concentrations. At low concentration (100 μM), Spm causes weak membrane hyperpolarization and transient H^+^ but not Ca^2+^ efflux. In contrast, higher (1 mM) Spm causes membrane depolarization in a ROS-independent manner (Pottosin et al., 2014). In contrast to Spm, other polyamines only trigger membrane depolarization at any given concentration (Pottosin et al., 2014). ROS derived from polyamine oxidation can also activate a variety of non-selective Ca^2+^-permeable channels leading to increases in [Ca^2+^]_cyt_ (Pei et al., 2000; Demidchik et al 2003 and 2007). Externally supplied polyamines can also trigger nitric oxide (NO) generation and intracellular Ca^2+^release through a pathway involving cGMP and cADPR (Neill et al., 2002). Besides, polyamines can also stimulate Ca^2+^ efflux by activation of plasma membrane Ca^2+^-ATPase activity (Pottosin et al., 2014). To note that plasma membrane Ca^2+^-ATPases ACA 8 (ARABIDOPSIS-AUTOINHIBITED Ca^2+^-ATPase 8) and ACA10 are found in complex with FLS2, and the double *aca8 aca10* mutant shows decreased flg22-induced Ca^2+^ and ROS burst (dit Frey et al., 2012). However, the identity of the Ca^2+^ channels and detailed mechanisms that mediate the Ca^2+^ signature triggered by Spm remain largely unknown.

To further investigate the effect of Spm and Put on PTI, we performed global gene expression analyses in the wild-type at 24 h of treatment with flg22, Spm, Put and combinations (Spm + flg22 and Put + flg22). Despite the inhibitory effect of Spm on the observed flg22-triggered ROS burst, transcriptional responses to flg22 were only partly compromised (**Figures 7 and 8**). Spm-treated plants were responsive to flg22 although *Pst* DC3000 resistance was partly compromised (**Figure 3A**). Both Put and Spm triggered the upregulation of genes related to cell wall biogenesis. However, the polyamines exhibited differences in the transcriptional responses of genes in primary metabolism, transcription factors, protein synthesis and degradation (**Figure 5**). In a previous report (Liu et al., 2020), we found that polyamines exhibited similar early transcriptional responses at 1 h of treatment. Differences between early and late responses were also observed in PAMPs such as flg22 or oligogalacturonides (OGs) probably due to hormonal and metabolic adaptation (Denoux et al., 2008). The flg22, (Spm+ flg22) and (Put + flg22) treatments also revealed that polyamines differentially reshape the transcriptional responses to PAMPs. In particular, Spm and less markedly Put, compromised transcriptional responses to flg22 related to ribosome biogenesis while they increased the expression of subsectors of flg22-responsive related to defense (**Figures 7, 8, S10 and S15**). Transcriptional potentiation of late PTI responses might be associated to delayed but sustained ROS production triggered by polyamines **(Figure S3)**. In addition, RBOHD activity has been shown to antagonize SA-induced pro-cell death signals in Arabidopsis (Torres et al., 2005), thus revealing a complex interrelationship between ROS and defense outputs. (Bach-Pages et al., 2020) found that the RNA-binding activity of eurkaryotic initiation factors, elongation factors and ribosomal proteins was inhibited in response to flg22. The upregulation in gene expression of ribosomal proteins caused by flg22 might be a compensatory mechanism to the PAMP-triggered inhibition of translation. In contrast, the polyamines Spd and Spm, and less strongly Put, are known to stimulate translation elongation and thus protein biosynthesis, which might compensate the inhibitory effect of flg22 on translation (Igarashi and Kashiwagi, 2015; Dever and Ivanov, 2018). Collectively, we conclude that polyamines differentially modulate PTI responses including Ca^2+^ signals and ROS production ultimately leading to changes in global transcriptional responses with an impact on defense against *P. syringae* in Arabidopsis.

## MATERIALS AND METHODS

### Plant materials

Seeds of the different genotypes were sown on a soil mixture containing peat moss (40%), vermiculite (50%) and perlite (10%), and stratified in the dark at 4°C for 2-3 days. Plants were grown at 20-22°C under 12-hour light/ 12-hour dark photoperiod cycles as described (Liu et al., 2020). Genotypes used in this work were obtained from the Eurasian Arabidopsis Stock Center (https://arabidopsis.info/) or previously described: *adc1-3, adc2-4* and *spms* (Alcázar et al., 2006; Cuevas et al., 2008; Liu et al., 2019), *eds1-2* (Feys et al., 2005), *pad4-1* (Glazebrook et al., 1997), *sid2-1* (Wildermuth et al., 2001), *npr1-1* (Cao et al., 1997), *fls2* (Heese et al., 2007), *rbohd* N663633 (SALK_109396C), *rbohd* N670541 (SALK_035391C), *rbohf* N657584 (SALK_034674C), *rbohd/f* N9558 (CS9558), *atao1* N672056 (SALK_127609C), *cuao1* N608014 (SALK_108014), *cuao2* N677606 (SALK_012167C), *cuaoα1* N661128 (SALK_125537C), *cuaoα2* N677690 (SALK_037584C), *cuaoδ* N686526 (SALK_094630C), *cuaoε1* N670103 (SALK_124509C), *cuaoε2* N730426 (GK-422D03.08), *cuaoγ2* N2054517 (GK-051A08.10), *pao1* N658095 (SALK_013026C), *pao2* N660420 (SALK_049456C), *pao3* N668943 (SALK_121288C)*, pao4* N653495 (SALK_133599C), *pao5* N679676 (SALK_053110C).

### ROS measurements

The detection and quantitation of reactive oxygen species (ROS) was performed by monitoring luminescence using a microplate luminometer (Luminoskan, Thermo Fisher). Leaf discs (0.5 cm diameter) were cut with a cork borer using fully expanded leaves of 5-week-old plants and incubated in 200 μL sterile water in 96-well plates during 24h. The water was then replaced with a solution containing 10 μg/ml horseradish peroxidase (HRP) (Merck), 100 μM of the luminol derivate L-012 (Wako Chemicals) and the different treatments at indicated concentrations. For pre-incubation assays, leaf discs were pretreated with Put (100 μM) or Spm (100 μM) 24 h before flg22 (1 μM) elicitation.

### Pathogen infection assays

Bacteria (*Pseudomonas syringae* pv. *tomato* DC3000, Pst DC3000) was inoculated in fully expanded leaves of 5-week-old plants by syringe-infiltration using a bacterial suspension (OD_600_ =0.005) in 10 mM MgCl_2_. *Pst* DC3000 colony forming units (CFU) per cm^2^ were determined at 72 h post-inoculation as described (Liu et al., 2020) using eight biological replicates per treatment and genotype.

### Pharmacological treatments

Leaf discs (0.5 cm diameter) were cut with a cork borer using fully expanded leaves of 5-week-old plants. The leaf discs were incubated in 200 μL sterile water in 96-well plates during 24h. Then, water was removed and the discs incubated for 3 h in a solution containing the different pharmacological treatments. Upon incubation, flg22 was added to 1 μM final concentration and ROS production monitored by luminescence as described above. The pharmacological treatments were as follows: 5 mM Dimethylthiourea (DMTU), 5 mM 2-Bromoethylamine hydrobromide (BEA), 20 μM Diphenyleneiodonium chloride (DPI), 1 mM reduced L-Glutathione (GSH), 2 mM EGTA, 1 mM LaCl_3_, 50 μM or 300 μM Cycloheximide (CHX), 20 μM Latrunculin B (Lat B) and 2.5 μM Brassinazole (BRZ). Upon flg22 elicitation, ROS production was monitored by luminescence detection as described above.

### DAB and Trypan blue staining

3,3’-diaminobenzidine (DAB) staining was performed by incubation of leaves in 3,3’-diaminobenzidine tetrahydrochloride (1 mg/mL, pH 3.8) overnight followed by de-staining in 100% ethanol for 3 h (Clarke, 2009). Trypan blue staining for cell death visualization was performed as previously described (Alcázar et al., 2009).

### Ca^2+^ measurements

A transgenic (Col-0) line expressing cytosolic apoaequorin was used for the quantitation of cytosolic Ca^2+^ (Knight et al., 1991). Leaf discs from 5-week-old plants expressing apoaequorin were incubated in 10 μM coelenterazine for 24 h in the dark in 96-well plates. Afterwards, the liquid was replaced with 100 μl ddH2O. Luminescence was continuously recorded during the different treatments using a microplate luminometer (Luminoskan, ThermoFisher Scientific). To calculate absolute cytoplasmic Ca^2+^ concentrations, the remaining aequorin in each microplate well was discharged by adding 100 μL of 2 M CaCl_2_ in 20% ethanol (Fricker et al., 1999; Maintz et al., 2014).

### RNA-seq gene expression analyses

Polyamine, flg22 and mock (water) treatments were performed in three biological replicates by leaf-infiltration of 5-week-old wild-type (Col-0) plants. Infiltrated leaves were collected and frozen in liquid nitrogen at 24 h of treatment for total RNA extraction. Total RNA was extracted using *TriZol* (Thermo Fisher) and further purified using RNeasy kit (Qiagen) according to manufacturer’s instructions. Total RNA was quantified in Qubit fluorometer (Thermo Fisher) and checked for purity and integrity in a *Bioanalyzer-2100* device (Agilent Technologies). RNA samples were further processed by the Beijing Genomics Institute (BGI) for library preparation and RNA sequencing using DNBSEQ. Libraries were prepared using the MGIEasy RNA Library Prep kit (MGI Tech) according to manufacturer’s instructions and each library was paired-end sequenced (2 x 100 bp) on DNBSEQ-G400 sequencers. Read mapping and expression analyses were performed using the *CLC Genomics Workbench 21 version 21.0.5* (Qiagen). Only significant expression differences (fold-change ≥ 2; Bonferroni corrected *p*-value ≤ 0.05) were considered. PCA, HCA and gene ontology analyses were performed using *CLC Genomics Workbench 12 version 21.0.5* (Qiagen) and Gene Ontology resource (GO; http://geneontology.org) using annotations from Araport11 (Cheng et al., 2017; Carbon et al., 2019). Pathway enrichment analyses were performed using PLANTCYC 15.0.1 (https://plantcyc.org/) and manual inspection of differentially expressed genes.

## Accession numbers

RNA-seq data has been deposited to ArrayExpress (https://www.ebi.ac.uk/arrayexpress/) under accession number X-XXXX-XXXX.

## ACKNOWLEDGEMENTS AND FUNDING

This work was funded by the BFU2017-87742-R grant of the Programa Estatal de Fomento de la Investigación Científica y Técnica de Excelencia (Ministerio de Economía y Competitividad, Spain), the Agencia Estatal de Investigación (AEI, Spain) MCIN/ AEI/10.13039/501100011033 and the Fondo Europeo de Desarrollo Regional (FEDER) “ERDF a way of making Europe”. Zhang Chi acknowledges support from the CSC (China Scholarship Council) for funding his doctoral fellowship. In memoriam of Prof. Antonio F. Tiburcio (1952-2021) for his many contributions to the research on polyamines in plants.

## AUTHOR CONTRIBUTIONS

C.Z., K.E.A and R.A. planned the experiments; C.Z and K.E.A. performed most of the experimental work; C.Z and R.A. analyzed the data; R.A. conceived the project and wrote the article with contributions from all authors.

## SUPPLEMENTAL FIGURES (Figure S1 – S20)

**Figure S1.** Effect of the Put and Spm cotreatment on flg22-elicited ROS burst. Leaf discs from 5-week-old wild-type plants were treated with flg22 (1 μM), and equal concentrations of Put and Spm (50 μM to 400 μM). Values represent the mean ± S.E. from at least twelve replicates per treatment and are expressed in relative light units (RLU).

**Figure S2**. Effect of Spm on flg22-elicited ROS burst in *eds1-2, pad4-1, sid2-1, npr1-1* and *fls2* negative control. Leaf discs from 5-week-old wild-type plants were treated with flg22 (1 μM), Spm (100 μM), Spm (100 μM) + flg22 (1 μM) or mock (water). Values represent the mean ± S.E. from at least twelve replicates per treatment and are expressed in relative light units (RLU).

**Figure S3**. ROS produced by Put and Spm treatment. Leaf discs from 5-week-old wild-type plants were incubated with different concentrations (100 μM to 800 μM) of Put, Spm and mock (water). Values represent the mean ± S.E. from at least twelve replicates per treatment and are expressed in relative light units (RLU).

**Figure S4**. Effect of Spm on flg22-elicited ROS burst in *rbohd* (N663633 and N670541), *rbohf* (N657584) and double *rbohd/f* (N9558) mutants. Leaf discs from 5-week-old wild-type plants and mutants were treated with flg22 (1μM), Spm (100 μM) or Spm (100 μM) + flg22 (1 μM). Values represent the mean ± S.E. from at least twelve replicates per treatment and are expressed in relative light units (RLU).

**Figure S5.** Analysis of *Pst* DC3000 disease resistance phenotypes in wild-type plants locally pretreated with different concentrations of Put (0 μM to 500 μM). Treatments were performed 24 h before *Pst* DC3000 infiltration (OD_600 nm_= 0.005). Bacterial numbers were assessed at 72 h post-inoculation and expressed as colony forming units (CFU) per cm^2^ leaf area. Values are the mean from at least eight biological replicates ±SD. Letters indicate values that are significantly different according to Tukey’s HSD test at *P* < 0.05.

**Figure S6. (A)** Flg22-elicited ROS and **(B)** effect of Spm on flg22-elicited ROS production in *CuAO* mutants (*atao1, cuao1, cuao2, cuaoα1, cuaoα2. cuaoδ, cuaoε1, cuao&2, cuao*γ2) and *PAO* mutants (*pao1, pao2, pao3, pao4* and *pao5*) in comparison to the wild-type. The total sum of RLU in each genotype was normalized to the total sum of RLU in the wild-type reference. Values represent the mean ± S.E. of the normalized values from at least twelve replicates per treatment. Letters indicate values that are significantly different according to Tukey’s HSD test at *P* < 0.05.

**Figure S7.** Effect of 2-bromoethylamine BEA (5 mM), diphenyleneiodonium chloride (DPI, 20 μM), dimethylthiourea (DMTU, 5 mM), reduced L-glutathione (GSH, 1 mM), EGTA (2 mM), LaCl_3_ (1 mM), cycloheximide (CHX, 300 μM) and Latrunculin B (Lat B, 20 μM) on Spm inhibition of flg22-triggered ROS burst in wild-type plants. Leaf discs from 5-week-old wildtype plants were pretreated with the different chemicals 3 h before Spm (100 μM), flg22 (1 μM) and Spm (100 μM) + flg22 (1 μM) elicitation. RLU counts were determined over time. Values represent the mean ± S.E. from at least twelve replicates per treatment. Letters indicate values that are significantly different according to Tukey’s HSD test at *P* < 0.05.

**Figure S8. (A)** Effect of brassinazole (BRZ, 2.5 μM) on Spm inhibition of flg22-triggered ROS burst in the wild-type. **(B,C)** Effect of cycloheximide (CHX, 50 μM) on Spm inhibition of flg22-triggered ROS burst in **(B)** wild-type and *rbohd* and **(C)** wild-type and *fls2*. Treatments were performed as described in Figure S7. Values represent the mean ± S.E. from at least twelve replicates per treatment and are expressed in relative light units (RLU). Letters indicate values that are significantly different according to Tukey’s HSD test at *P* < 0.05.

**Figure S9. (A)** 3,3’-diaminobenzidine (DAB) staining of wild-type and *rbohd* mutants infiltrated with Spm (100 μM), methyl viologen (100 μM) or both (100 μM Spm + 100 μM MV). Staining was performed at 24 h of treatment. **(B)** Trypan blue staining of wild-type and *rbohd* leaves infiltrated as in (A).

**Figure S10.** Gene expression values and correlation analyses of wild-type (Col-0) plants treated with flg22 (1 μM), Spm (100 μM) and Spm (100 μM) + flg22 (1 μM). **(A)** Genes only significantly deregulated by flg22 treatment. **(B)** Common genes deregulated by flg22 and (Spm + flg22) treatments. **(C)** Genes only deregulated by (Spm + flg22) treatment. **(D)** Genes only deregulated by Spm treatment. Expression values (Log2) are relative to the mock (H_2_O). The mean expression ± S.E of upregulated and downregulated genes is shown for each treatment. Asterisks indicate significant differences according to Wilcoxon signed-rank test (***p<0.001). **(E)** Expression correlation between flg22, Spm and Spm+flg22 treatments.

**Figure S11.** Molecular function categorization and metabolic pathway enrichment analysis of genes only deregulated by flg22 compared to Spm and (Spm + flg22) treatments in the wild-type. Bars indicate the mean expression ± S.E of each category.

**Figure S12.** Molecular function categorization and metabolic pathway enrichment analysis of genes commonly deregulated in flg22 and (Spm + flg22) treatments in the wild-type. Bars indicate the mean expression ± S.E of each category.

**Figure S13.** Molecular function categorization and metabolic pathway enrichment analysis of genes only differentially expressed in (Spm + flg22) compared to flg22 and Spm treatments. Bars indicate the mean expression ± S.E in each category.

**Figure S14.** Molecular function categorization and metabolic pathway enrichment analysis of genes only differentially expressed in Spm treatment compared to flg22 and (Spm + flg22) treatments. Bars indicate the mean expression ± S.E of each category.

**Figure S15.** Gene expression values and correlation analyses in wild-type (Col-0) plants treated with flg22 (1 μM), Put (100 μM) and Put (100 μM) + flg22 (1 μM). **(A)** Genes only significantly deregulated by flg22 treatment. **(B)** Common genes deregulated by flg22 and (Put + flg22) treatments. **(C)** Genes only deregulated by (Put + flg22) treatment. **(D)** Genes only deregulated by Put treatment. Expression values (Log2) are relative to the mock (H_2_O). The mean expression ± S.E of upregulated and downregulated genes is shown for each treatment. Asterisks indicate significant differences according to Wilcoxon signed-rank test (***p<0.001). **(E)** Expression correlation between flg22, Put and Put+flg22 treatments.

**Figure S16.** Molecular function categorization and metabolic pathway enrichment analysis of genes only deregulated by flg22 compared to Put and (Put + flg22) treatments. Bars indicate the mean expression ± S.E of each category.

**Figure S17.** Molecular function categorization and metabolic pathway enrichment analysis of genes commonly deregulated in flg22 and (Put + flg22) treatments in the wild-type. Bars indicate the mean expression ± S.E of each category.

**Figure S18.** Molecular function categorization and metabolic pathway enrichment analysis of genes only deregulated in (Put + flg22) compared to flg22 and Put treatments. Bars indicate the mean expression ± S.E of each category.

**Figure S19.** Molecular function categorization and metabolic pathway enrichment analysis of genes only deregulated by Put compared to flg22 and (Put + flg22) treatments. Bars indicate the mean expression ± S.E of each category.

**Figure S20.** Effect of thermospermine (tSpm, 100 μM) and spermidine (Spd, 100 μM) on flg22-elicited ROS burst in the wild-type (Col-0). Leaf discs from 5-week-old plants were treated with flg22 (1 μM), tSpm (100 μM), Spd (100 μM), tSpm (100 μM) + flg22 (1 μM), Spd (100 μM) + flg22 (1 μM) or mock (water). Values represent the mean ± S.E. from at least twelve replicates per treatment and are expressed in relative light units (RLU).

## SUPPLEMENTAL TABLES

**<u>Tables S1.1 to S1.8.</u>** Genes differentially expressed at 24 h of Put (100 μM) and Spm (100 μM) treatments in the wild-type.

**Table S1.1.** List of 554 differentially expressed genes at 24 h of Put (100 μM) treatment.

**Table S1.2.** List of 368 differentially expressed genes at 24 h of Spm (100 μM) treatment.

**Table S1.3.** List of 396 genes only deregulated by Put (100 μM).

**Table S1.4.** List of 210 genes only deregulated by Spm (100 μM).

**Table S1.5.** List of 158 common genes which are differentially expressed in Spm (100 μM) and Put (100 μM) treatments.

**Table S1.6.** Gene ontology (GO) analysis of genes in Table S1.3

**Table S1.7.** Gene ontology (GO) analysis of genes in Table S1.4

**Table S1.8.** Gene ontology (GO) analysis of genes in Table S1.5

**<u>Tables S2.1 to S2.19.</u>** Comparison of genes which are differentially expressed at 24 h of treatment with flg22 (1 μM), Spm (100 μM) and Spm (100 μM) + flg22 (1 μM) in the wild-type.

**Table S2.1.** List of 1415 differentially expressed genes at 24 h of flg22 (1 μM) treatment.

**Table S2.2.** Gene ontology analysis of genes deregulated in Table S2.1

**Table S2.3.** List of 462 flg22-only genes that show significant expression differences at 24 h of flg22 (1 μM) treatment.

**Table S2.4.** Gene ontology analysis of genes deregulated in Table S2.3

**Table S2.5.** Pathway enrichment analysis of genes deregulated in Table S2.3

**Table S2.6.** List of 1534 differentially expressed genes at 24 h of (100 μM Spm + 1 μM flg22) treatment.

**Table S2.7.** Gene ontology analysis of genes deregulated in Table S2.6

**Table S2.8.** List of 868 common genes which are differentially expressed in flg22 (1 μM) and (100 μM Spm + 1 μM flg22) treatments.

**Table S2.9.** Gene ontology analysis of genes deregulated in Table S2.8

**Table S2.10.** Pathway enrichment analysis of genes deregulated in Table S2.8

**Table S2.11.** List of 543 (Spm + flg22)- only genes that show significant differences in expression at 24 h of (100 μM Spm + 1 μM flg22) treatment.

**Table S2.12.** Gene ontology analysis of genes deregulated in Table S2.11

**Table S2.13.** Pathway enrichment analysis of genes deregulated in Table S2.11

**Table S2.14.** List of 220 (Spm-only) genes that show significant differences in expression at 24 h of Spm (100 μM) treatment.

**Table S2.15.** Gene ontology analysis of genes deregulated in Table S2.14

**Table S2.16.** List of 25 common genes which are differentially expressed in flg22 (1 μM) and Spm (100 μM) treatments.

**Table S2.17.** List of 63 common genes which are differentially expressed in Spm (100 μM) and (100 μM Spm + 1 μM flg22) treatments.

**Table S2.18.** List of 60 common genes which are differentially expressed in Spm (100 μM), flg22 (1 μM) and (100 μM Spm + 1 μM flg22) treatments.

**Table S2.19.** Gene ontology analysis of genes deregulated in Table S2.18

**<u>Tables S3.1 to S3.19.</u>** Comparison of genes which are differentially expressed at 24 h of treatment with flg22 (1 μM), Put (100 μM) and Put (100 μM) + flg22 (1μM) in the wild-type.

**Table S3.1.** List of 259 flg22-only genes that show significant expression differences at 24 h of flg22 (1 μM) treatment.

**Table S3.2.** Gene ontology analysis of genes deregulated in Table S3.1

**Table S3.3.** Pathway enrichment analysis of genes deregulated in Table S3.1

**Table S3.4.** List of 1395 differentially expressed genes at 24 h of (100 μM Put + 1 μM flg22) treatment.

**Table S3.5.** Gene ontology analysis of genes deregulated in Table S3.4

**Table S3.6.** List of 902 common genes which are differentially expressed in flg22 (1 μM) and (100 μM Put + 1 μM flg22) treatments.

**Table S3.7.** Gene ontology analysis of genes deregulated in Table S3.6

**Table S3.8.** Pathway enrichment analysis of genes deregulated in Table S3.6

**Table S3.9.** List of 300 (Put + flg22)-only genes that show significant differences in expression at 24 h of (100 μM Put + 1 μM flg22) treatment.

**Table S3.10.** Gene ontology analysis of genes deregulated in Table S3.9

**Table S3.11.** Pathway enrichment analysis of genes deregulated in Table S3.9

**Table S3.12.** List of 270 (Put-only) genes that show significant differences in expression at 24 h of Put (100 μM) treatment.

**Table S3.13.** Gene ontology analysis of genes deregulated in Table S3.12

**Table S3.14.** Pathway enrichment analysis of genes deregulated in Table S3.12

**Table S3.15.** List of 91 common genes which are differentially expressed in the treatments with Put (100 μM) and flg22 (1 μM).

**Table S3.16.** Gene ontology analysis of genes deregulated in Table S3.15

**Table S3.17.** List of 163 common genes which are differentially expressed in the treatments with Put (100 μM), flg22 (1 μM) and (100 μM Put + 1 μM flg22).

**Table S3.18.** Gene ontology analysis of genes deregulated in Table S3.17

**Table S3.19.** List of 30 common genes which are differentially expressed in the treatments with Put (100 μM) and (100 μM Put + 1 μM flg22).

